# Spatiotemporal Dynamics of Reward and Punishment Effects Induced by Associative Learning

**DOI:** 10.1101/349712

**Authors:** Huan Wang, Killian Kleffner, Patrick L. Carolan, Mario Liotti

## Abstract

While reward associative learning has been studied extensively across different species, punishment avoidance learning has received far less attention. Of particular interest is how the two types of learning change perceptual processing of the learned stimuli. We designed a task that required participants to learn the association of emotionally neutral images with reward, punishment, and no incentive value outcomes through trial-and-error. During learning, participants received monetary reward, neutral outcomes or avoided punishment by correctly identifying corresponding images. Results showed an early bias in favor of learning reward associations, in the form of higher accuracy and fewer trials needed to reach learning criterion. We subsequently assessed electrophysiological learning effects with a task in which participants viewed the stimuli with no feedback or reinforcement. Critically, we found modulation of two early event-related potential components for reward images: the frontocentral P2 (170 – 230 ms) and the anterior N2/Early Anterior Positivity (N2/EAP; 210 – 310 ms). We suggest that reward associations may change stimuli detection and incentive salience as indexed by P2 and N2/EAP. We also reported, on an exploratory basis, a late negativity with frontopolar distribution enhanced by punishment images.

## 1. Introduction

We remember not only individuals, objects, or situations from the past, but also their associations with pleasant or unpleasant outcomes. Such associations dramatically bias our decisions towards profitable outcomes [1-3]. Animal research suggests that neutral cues become highly attractive when they are paired with food reward, with rats attempting to consume the inedible cue items [4, 5]. This is because pairing with a food reward confers incentive salience to the conditioned stimulus, a mechanism dependent on the dopaminergic mesolimbic system, particularly the ventral striatum [6]. Human neuroimaging studies further suggest that the orbitofrontal cortex (OFC) is also involved in reward processing [7-11], which has been proposed to store higher-order cognitive representations of the incentive salience [12].

In contrast, while there is evidence showing neural correlates of punishment avoidance learning in humans [13-16], the mechanism through which avoiding punishment could influence the mental representation of cues and choices is not fully established. One account proposes that avoiding punishment itself can be rewarding, with the absence of an aversive outcome serving as a positive reinforcer to bias choice [14]. However, a lesion study found that damage to anterior insula and dorsal striatum selectively impaired punishment avoidance but not reward learning [15], suggesting some dissociations underlying the two types of learning. Nevertheless, at least in some cases, the OFC seems to contribute to both reward and punishment avoidance learning, which has led to the suggestion that OFC encodes an unsigned value representation for both reward and punishment cues [17, 18]. Yet, how punishment avoidance learning could change the mental representations of the predictive cues is not well understood.

Utilizing the measurement benefits of human electroencephalography (EEG) may provide another powerful way to study associative learning, as it provides a temporal characterization of exactly ‘when’ different neural processes are activated [19]. Using event-related potentials (ERPs), a human reward conditioning study found an early frontocentral (P2, 180 -280 ms) component that was larger for reward than neutral cues [20]. Participants also had an enhanced P2 when anticipating monetary reward with improved task performance on reward-cued trials. These results suggest that P2 modulations may reflect initial attentional capture towards reward cues [21-23]. Behavioral results have provided further support for such interpretation, showing that reward cues indeed attract attention and influence task performance, even when they are task-irrelevant [24-26].

Similar to P2 effects, another ERP component with frontal scalp distribution has been found to be modulated when participants anticipate monetary reward and prepare for subsequent responses (anterior N2, 180-300 ms; [21, 23]). Similar changes in N2 amplitude have also been reported when emotional stimuli serve as distractors to capture attention [27, 28]. We have previously reported a component as modulations of the anterior N2, called the Early Anterior Positivity (EAP; [29, 30]) using task-irrelevant but personally salient stimuli, such as when marijuana users viewed images of drug-related content and when chocolate cravers viewed images depicting chocolate [31, 32]. Additionally, consistent with previous fMRI results (e.g., [7]), source localization of the anterior N2/EAP showed a main source in OFC [31-33], making this another potential candidate for an electrophysiological index of the incentive salience representation.

Classical conditioning studies using punishment (e.g., aversive odors or electric shock) have found mainly early visual modulations at occipital sites [34, 35]. One study found an early prefrontal modulation to aversively conditioned cues, which was proposed to be a part of the circuit that detected potential threats in the environment [36]. However, none of the studies focused on punishment avoidance learning, or compared punishment avoidance learning to reward learning. One recent ERP study has incorporated elements of both monetary gain and loss in the course of trial-and-error learning. In this task, participants learned to identify novel characters that were associated with monetary gain, loss, or neutral outcomes. Characters were presented in pairs and participants learned to pick the rewarded characters to achieve maximum payoff. In the testing phase, scalp EEG was recorded while participants made old/new judgments on learned versus novel characters. Modulations of early occipital and late parietal components were found for the rewarded characters, while no changes were found frontally or for the punishment characters [37]. Because only limited behavioral results were reported from the learning phase, it was difficult to interpret how reward learning was related to the reward ERP effects. Similarly, punishment associations were actually established through incurring monetary loss rather than avoiding such loss, which offered limited insight on punishment avoidance learning and its subsequent effects on perception.

To address the limitations of the previous studies, we designed a paradigm involving an associative learning task with both monetary reward, punishment and a neutral control condition. Critically, unlike previous studies in which the task was terminated once the learning criterion was achieved [37] or incorporated a limited number of learning trials (e.g., [38]), we set the period of learning to be sufficiently long (30 minutes) allowing participants to continue the task for the full duration even after reaching the learning criterion. This generated enough trials to allow for a thorough learning analysis. Moreover, three groups of cues were assigned with three different response buttons and strategically pressing only one button produced no long-term benefit. Following the associative learning task, all cues were presented in the context of an *implicit* task in which participants were instructed to attend to the physical characteristics of the cues. No feedback or monetary contingency was provided for this task. We also examined how behavioral data of learning were related to the subsequent perceptual processing of the cues.

Our first hypothesis was that an ERP modulation of the frontocentral P2 would be observed for reward cues as an index of early attentional capture. The frontal P2 is consistently reported in studies of reward anticipation [22, 23] and reward conditioning [20]. This is consistent with the notion that cues with reward incentive salience served as a “motivational magnet” attracting attention towards it [4, 6]. Secondly, we predicted that another potential ERP index of incentive salience often associated with emotional stimuli, the N2/EAP, would be enhanced for reward cues [29, 31, 39].

The last goal of the current report was to explore distinctive ERPs elicited during perceptual processing of punishment associated images following avoidance learning. No previous ERP component has been reported relating to punishment associations. Evidence from fMRI literature suggests that the OFC and anterior insula, in conjunction with the dorsal striatum, are implicated in avoidance learning and access to punishment representations [10, 14, 15]. An electrophysiological counterpart of activity generated by these brain regions is likely to yield effects over anterior frontal scalp locations, which we looked for in our data, without a clear prediction of the exact timing of such activity. We resolved to conduct an exploratory analysis on a putative punishment-related frontal component that could shed light on future research on punishment association and avoidance learning.

## 2. Methods

### 2.1. Participants

Forty participants (26 female, 32 right-handed, age mean = 20.4, range = 17-33) took part in the experiment in exchange for course credits. All reported having normal or correct-to-normal vision, and none was color-blind. The Simon Fraser University Research Ethics Board approved all components of the current study.

### 2.2. Procedure

After providing informed consent, participants were prepared for EEG recording while they completed a demographic questionnaire. They were then transferred to a sound-attenuated room and completed the learning task, followed by the recording task. Immediately after, participants completed a subjective rating task and an affective priming task, the data of which will not be a main focus of the current report. Sessions were 2-hours long, with all tasks completed on the same day.

### 2.2.1. Associative learning task

Participants viewed a serial presentation of stimuli displayed on an LCD monitor against a black background. They were informed that each stimulus belonged to one of three preassigned groups: reward, punishment, or neutral. Stimuli consisted of 18 pictures depicting mundane objects or abstract scenes (e.g., a plant, a book, a spoon, colored shapes etc.) selected from the International Affective Picture System (IAPS; [40])^1^. Stimuli preassigned to the three groups had normative mean valence ratings of 4.85 (0.21), 5.03 (0.24), 5.12 (0.18), and arousal ratings of 2.52 (0.47), 2.74 (0.77), 2.50 (0.39). One-way analyses of variance (ANOVA) indicated that these ratings were not significantly different from one another (*ps* > .14). The three preassigned groups were counterbalanced across participants to minimize perceptual differences or subjective affective responses.

Stimuli were presented in pseudo-random order, with the restriction that all groups had to be sampled from once before the next iteration, and repeated for a total of 30 minutes. Participants were instructed to indicate on each trial the category they believed the stimulus belonged to by pressing the left, down, or right arrow key on a keyboard. Stimuli remained on screen until the participant responded and were followed by a feedback stimulus showing both the actual and alternative outcomes for 1000 ms. Correct responses in the reward condition earned 5 points, incorrect responses in the punishment condition lost 5 points, and no point was assigned otherwise (Fig. 1A). Participants were asked to pay equal attention to all stimuli, independent of their incentive value. All points earned in the task were paid in Canadian dollars (CAD) with a conversion rate of 200:1 *(M* = 4 CAD). No EEG was recorded during learning due to concerns with eye blinks as well as distraction from learning.

### 2.2.2. Recording task

The 18 stimuli from the associative learning task were presented again in blocks of 20 trials. Two images were selected to be fillers and required responses from participants. Stimuli were presented for 500 ms, followed by a randomly jittered inter-trial interval of 1300 to 2000 ms. A central fixation cross was displayed throughout the task. Participants were instructed to respond to stimuli that they saw repeated a second time in that block, by pressing the key “1” on the number pad of a keyboard with their right-hand index finger (Fig. 1B). Images selected as filler trials and their locations of occurrence were randomized between blocks but consistent across participants. Filler trials and response errors (i.e., false alarms to non-fillers) were dropped from ERP analyses. Participants completed 20 blocks of the task, separated by short breaks to rest.

**Fig. 1.**
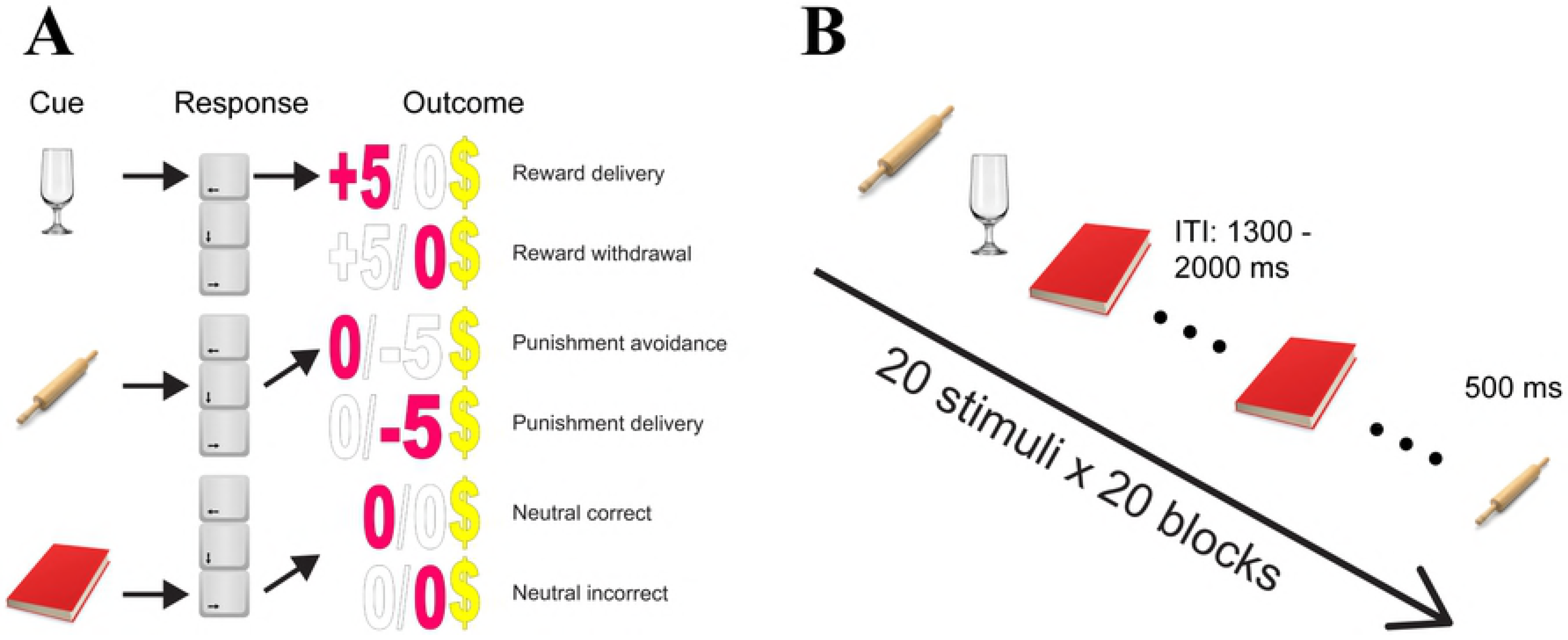
(A) *Associative learning task*: participants learned from trial-and-error which stimuli were from reward, punishment or neutral group. Each group of cues was preassigned with one button (left, down or right arrow key). Participants gained five points by correctly responding to the reward cues whereas they lost five points by incorrectly responding to the punishment cues. Zero point was given to all other cases. All points were reimbursed with Canadian dollars with a ratio of 200:1. Learning was self-paced, and lasted for 30 minutes for all participants. Feedback including the received and the alternative outcomes was shown for 1000 ms after each response. (B) *Recording task*: each block had 20 trials that featured the 18 previously learned stimuli from the associative learning task, two of which were repeated within each block as fillers. Participants were asked to respond to the fillers. All stimuli were shown for 500 ms, with an inter-trial interval (ITI) jittered between 1300 to 2000 ms. A fixation cross was superimposed to the center of the stimuli across the task. EEG was recorded during this task and participants were given breaks between blocks to rest and stretch. Note stimuli in the figure were for illustration only and see methods (2.2.1.) for IAPS reference numbers.

## 2.3. EEG recording and processing

Continuous EEG was acquired during the recording task from 64 active Ag/AgCl electrodes based on the standard 10-10 montage (BioSemi Active Two system, Amsterdam, NL). Two channels the Common Mode Sense (CMS) and Driven Right Leg (DRL) formed a feedback loop that served as the ground. EEG was referenced to CMS during online recording without any filter. Six additional channels were applied: one at each lateral canthus (for horizontal eye movements), one below each eye (for vertical eye movements and blinks) and one over each mastoid bone. Continuous EEG was digitized at a sampling rate of 512 Hz. Semi-automatic artifact rejection was implemented using BESA with a 120 μV cut-off from 200 ms before to 800 ms after the stimulus onset. All trials were also visually inspected for additional artifacts. On average, a proportion of .34 (.19), .33 (.19) and .33 (.19) of the trials were rejected for the reward, punishment and neutral condition respectively, with no significant difference among them (*p* > .25). All signals were filtered offline with a high-pass filter of 0.01 Hz (zero phase, 12 dB/octave slope; BESA 5.3, MEGIS Software GmbH, Gräfelfing, Germany) and a low-pass filter of 30 Hz (zero phase, 12 dB/octave slope; FieldTrip, [41]). Bad channels were interpolated using spherical spline method implemented in FieldTrip [41]. Four participants were dropped from ERP analysis due to noisy mastoid channel data, technical difficulties or poor overall data quality producing less than 30 trials per condition, resulting in 36 participants as our final sample. ERPs were time-locked to stimulus onset, aligned with a 200 ms prestimulus baseline, and re-referenced to average mastoids.

## 2.4. ERP components of interest

To allow for the selection of the regions of interest (ROIs), and the latency of the effects to be independent (orthogonal) to the conditions of interest, ROIs and time windows of the ERPs were chosen on a-priori predictions based on published data and upon visual inspection of the grand-average ERPs collapsed across conditions [42]. First, the anterior P2 peaked around 200 ms on the collapsed grand-average ERPs; A ROI including frontocentral electrodes (FC1, FCz, FC2 and Cz) and a time window between 170 and 230 ms were chosen based on previous literature reporting salience-related P2 effects [20, 21, 23, 42].

Next, the N2 peaked at 260 ms; a left anterior frontal ROI was selected (Fp1, AF7, AF3, F7, F5 and F3), based on previous literature highlighting the N2 as modulated by biological salience [27, 33], as well as results from our lab showing a left-lateralized N2/EAP effect for personally salient appetitive stimuli [31, 32]. Time window was set between 210 and 310 ms.

Finally, we observed a later frontopolar punishment-related ERP effect evident only upon inspection of the condition-specific grand-average ERPs, in the form of a sustained larger *negativity* to punishment than reward and neutral stimuli, spanning from 350 ms to about 800 ms (peaking around 450 ms; named Frontopolar Negativity, or FN; Fig. 2). In the absence of prior studies in the literature reporting such effect, the orthogonality requirements were not met [42]. An exploratory analysis was carried out on such punishment-related frontal effect, employing an ROI including sites Fp1, Fp2, AF3, AF4, AF7 and AF8 (the electrode locations of larger amplitude), and a time window of 350 to 650 ms. For all three ERP time windows (P2, N2/EAP, FN) mean amplitudes were extracted for statistical analyses.

**Fig. 2.**
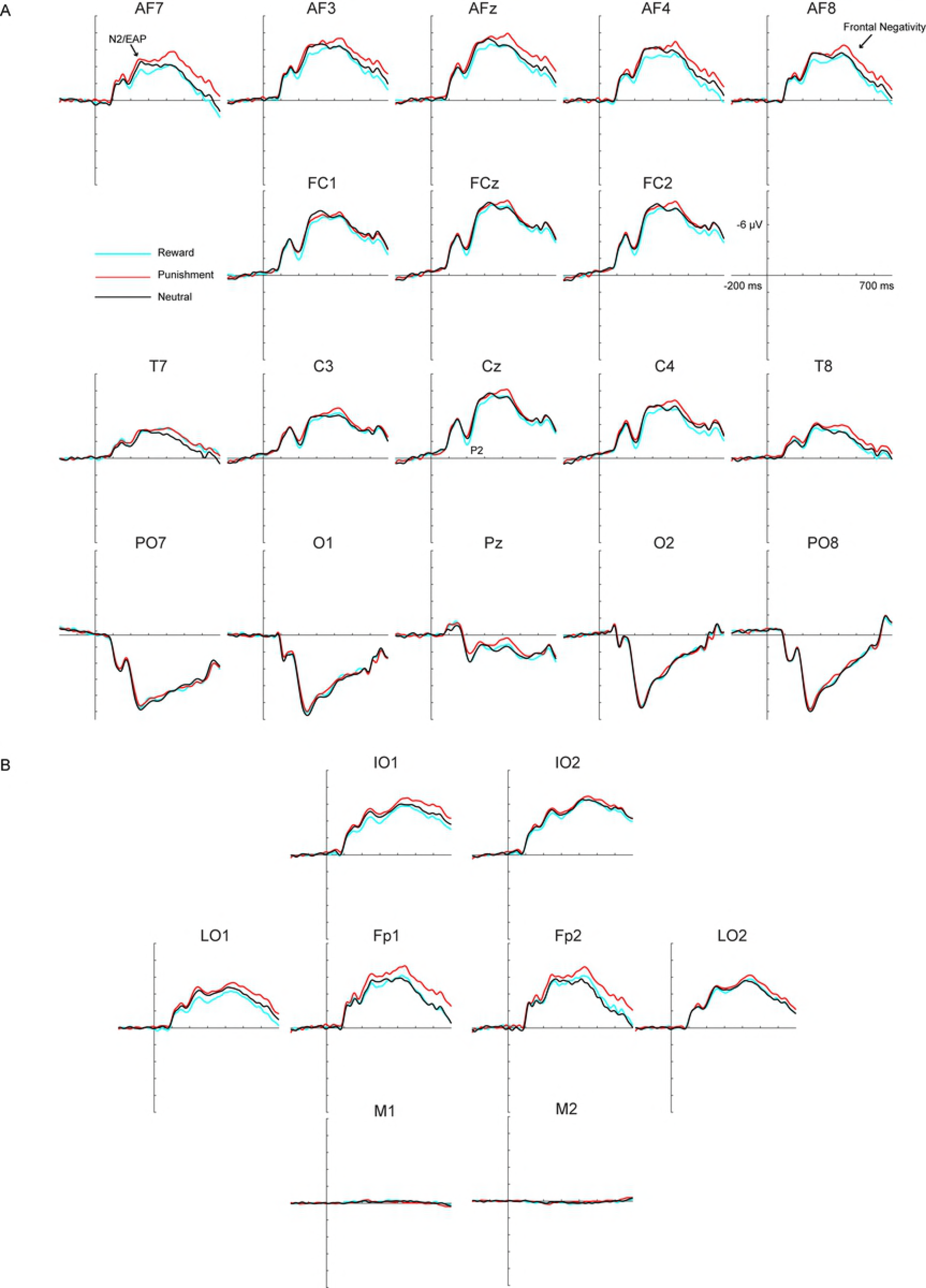
(A) Grand-averaged ERP waveforms for reward, punishment and neutral stimuli (N = 36). Waveforms reflect a selected array of 18 electrodes. Positive voltages are shown below the abscissa. The epoch is shown from -200 ms to 700 ms and each tick on the abscissa represents 100 ms. (B) Grand-averaged waveforms for all external electrodes and electrodes above the eyes. IO1-2: left and right infraorbital (below the eyes), LO1-2: left and right lateral ocular (beside the eyes), M1-M2: left and right mastoid, Fp1-2: left and right frontopolar (above the eyes). Note how the peak corresponding to the FN over frontopolar sites does not invert polarity below the eyes, or between left and right lateral ocular channels.

## 2.5. Analysis

### 2.5.1. Associative learning task

For each participant and condition (reward, punishment, neutral) mean accuracy and median reaction time (RT) were computed. Additionally, we estimated the learning points for each condition by identifying the first trials in which participants executed at least six consecutive correct responses. We chose this criterion because there were six pictures from each condition and correctly responding to all stimuli in a particular condition was the evidence of learning. RT and accuracy data were subgrouped into pre-learning and post-learning phases. RT and accuracy data that were more than 2 SDs from the means were considered outliers and were excluded from analyses. Cases where participants did not meet the learning criterion for a condition were dropped on an individual basis. After exclusion the final sample contained 29 subjects for RT and 27 for accuracy analyses. Finally, we collected the total number of button presses for left-arrow, down-arrow and right-arrow keys across learning.

RT and accuracy data were entered into two-way ANOVA with condition and learning (pre-learning, post-learning) as within-subjects factors. Learning point and button-press data were entered into one-way ANOVA with condition as within-subjects factor.

### 2.5.2. Recording task

For each participant and condition mean accuracy was computed separately for hits as well as false alarms. Sensitivity index d-Prime was calculated by subtracting z-transformed false alarm score from hits z-score. For each participant, ERP mean amplitudes (P2, N2/EAP, FN windows) were entered into one-way ANOVAs with condition as within-subject factor. For all ANOVAs (behavioral and ERP data), *post hoc* t-tests used the Bonferroni family-wise error correction to assess significant main effects and interactions. Alpha level was set to .05 and the Greenhouse-Geisser correction was applied when the assumption of sphericity was violated.

## 3. Results

### 3.1. Behavioral effects

#### 3.1.1. Associative learning task

The ANOVA of the learning point estimation returned a main effect of condition (*F* (1.6, 57.60) = 11.89, *p* < .001, ƞ^2^_p_ = .25), with reward cues being learned *faster* than punishment and neutral cues (*t* (36) = -2.85, *p* = .022; *t* (36) = -4.25, *p* < .001, respectively; see Table 1). There was also a trend for faster learning of punishment over neutral associations (*t* (36) = -2.32, *p* = .078). The median number of trials participants saw per condition was 245, with a range of 142 to 336 trials.

**Table 1.**
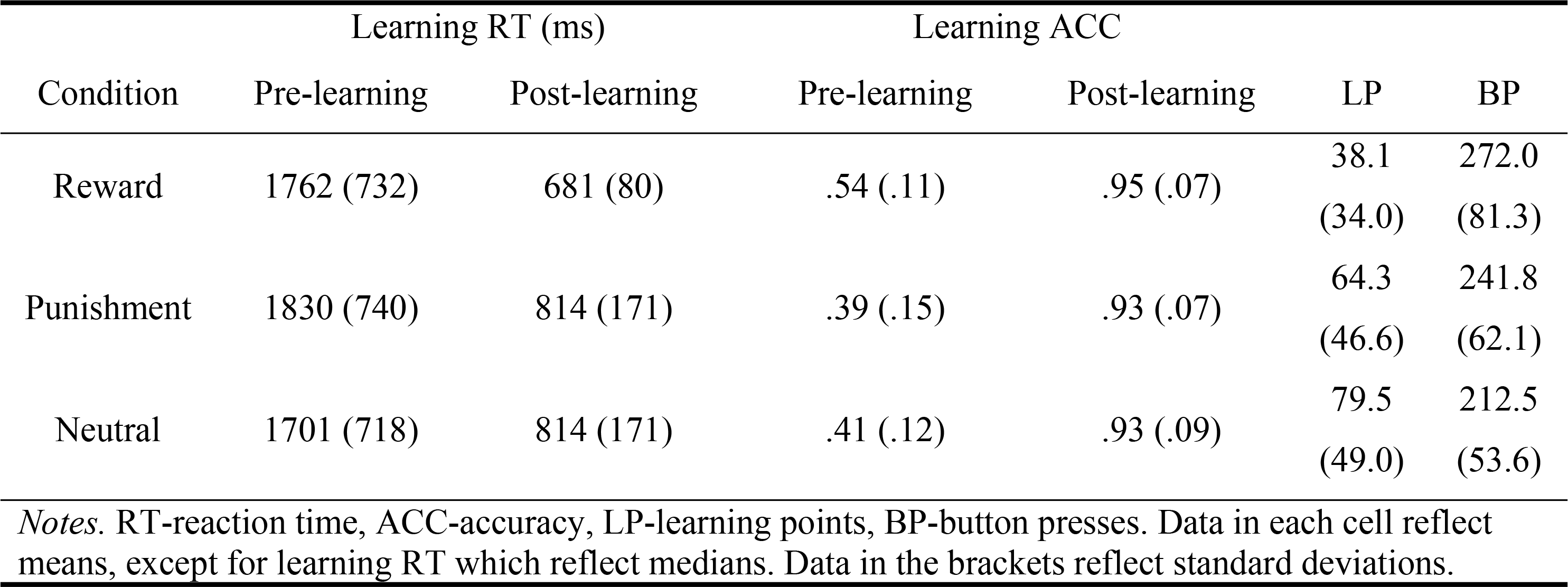
Behavioral results from associative learning.

| Condition | Learning RT (ms) |  | Learning ACC |  | LP | BP |
| --- | --- | --- | --- | --- | --- | --- |
|  | Pre-learning | Post-learning | Pre-learning | Post-learning |  |  |
| Reward | 1762 (732) | 681 (80) | .54 (.11) | .95 (.07) | 38.1<br>(34.0) | 272.0<br>(81.3) |
| Punishment | 1830 (740) | 814 (171) | .39 (.15) | .93 (.07) | 64.3<br>(46.6) | 241.8<br>(62.1) |
| Neutral | 1701 (718) | 814 (171) | .41 (.12) | .93 (.09) | 79.5<br>(49.0) | 212.5<br>(53.6) |

*Notes.* RT-reaction time, ACC-accuracy, LP-learning points, BP-button presses. Data in each cell reflect means, except for learning RT which reflect medians. Data in the brackets reflect standard deviations.

Button presses also showed a main effect of condition (*F* (2, 78) = 10.61, *p* < .001, ƞ^2^_p_ = .21), with reward and punishment buttons being pressed more than neutral buttons (*t* (39) = 4.25, *p* < .001; *t* (39) = 2.67, *p* = .033, respectively; Table 1).

Accuracy analysis showed a significant condition x learning two-way interaction (*F* (2, 52) = 7.00, *p* = .002, ƞ^2^_p_ = .21). It was explained by a main effect of condition *before* learning (*F* (2, 52) = 11.31, *p* < .001, ƞ^2^_p_ = .30; reward > punishment, *t* (26) = 4.09, *p* = .001, reward > neutral, *t* (35) = 3.47, *p* = .006).

In contrast, only trend level accurary difference was detected after learning (*F* (2, 52) = 3.29, *p* = .045, ƞ^2^_p_ = .11; reward > punishment, *t* (26) = 2.47, *p* = .061, reward > neutral, *t* (35) = 1.94, *p* = .19). These results suggested that participants showed an early learning bias towards reward cues, making more attempts towards receiving rewards.

The two-way ANOVA on split RT only returned a main effect of learning (*F* (1, 28) = 71.4, *p* < .001, ƞ^2^_p_ = .72), with faster RT after (*M* = 769.7 ms, *SD* = 23.3) than before learning (*M* = 1764.2 ms, *SD* = 119.2). No interaction or main effect of condition was observed (*ps* > .22).

#### 3.1.2. Recording task

No main effects were found for hit or false alarm rates (*p* > .26). Similarly, no d-prime difference was detected (*p* = .37; reward *M* = 1.49, *SD* = .67, punishment *M* = 1.31 *SD* =1.05, neutral *M* = 1.42, *SD* = .78). Collapsing across conditions, the data showed that participants followed the instructions well (false alarm *M* = .032, *SD* = .037) and remained engaged in the task (hit *M* = .54, *SD* = .19).

## 3.2. ERP effects

### 3.2.1. P2 mean amplitude (170-230 ms)

P2 amplitude showed a condition dependent difference (*F* (2, 70) = 3.23, *p* = .046, ƞ^2^_p_ = .084), with reward trials (*M* = -2.66 μV, *SD* = 4.11) having greater amplitude than punishment trials (*M* = -3.44 μV, *SD* = 4.45; *t* (35) = 2.70, *p* = .032, *d* = 0.18). The other two comparisons did not reach significance (reward vs. neutral, t (35) = 1.91, *p* = .19; punishment vs. neutral, t (35) = -.56, *p* = 1.0).

### 3.2.2. N2/EAP mean amplitude (210-310 ms)

Similarly, for the N2/EAP amplitude there was a main effect of condition (*F* (2, 70) = 3.63, *p* = .032, ƞ^2^_p_ = .094), due to a significant difference between the reward (*M* = -4.39 μV, *SD* = 3.72) and punishment conditions (*M* = -5.35 μV, *SD* = 3.56; *t* (35) = 2.88, *p* = .020, *d* = 0.27). The other two comparisons did not reach significance (reward vs. neutral, t (35) = 1.71, *p* = .29; punishment vs. neutral, t (35) = -.88, *p* = 1.0).

### 3.2.3. FN mean amplitude (350-650 ms)

Finally, we found a main effect of condition for FN (*F* (2, 70) = 4.58, *p* = .014, ƞ^2^_p_ = .12), as a result of difference between the punishment (*M* = -5.12 μV, *SD* = 4.83) and reward conditions (*M* = -3.78 μV, *SD* = 4.50; *t* (35) = -2.65, *p* = .036, *d* = 0.29), along with a trend level difference between punishment and neutral condition (*M* = -3.94 μV, *SD* = 3.93; *t* (35) = -2.26, *p* = .091, *d* = 0.27). Bear in mind, however, that the FN analysis was exploratory in nature and should be interpreted with caution.

### 3.3. Correlations to behavioral effects

Correlational analyses were conducted using Spearman’s rank-order test to account for significant outliers in our data. In light of our accuracy results, we looked at how pre-learning accuracy difference was related to P2 and N2/EAP difference waves of the significant contrasts (reward minus punishment). No difference was found (*p*s > .67).

However, we further explored more informative measures looking at how specific incorrect choices for each condition may impact learning. For example, when participants made errors for reward cues, they would make either punishment or neutral judgments. We speculated that making punishment judgments to reward cues might create a larger impact on learning and representations of the reward cues than making a neutral judgment. The same logic would apply to reward judgments towards punishment cues. Therefore, we calculated the difference score between the ratios of punishment errors on reward trials (i.e., updating reward associations) and reward errors on punishment trials. A higher value would suggest a stronger bias towards reward over punishment learning. Furthermore, with the inclusion of neutral cues, we calculated the difference between ratios of reward and punishment errors for neutral cues. This measure may suggest individual differences in decision under uncertainty independent of reward or punishment learning, given that errors on neutral trials only informed learning of neutral associations. We further controlled the general tendency towards choosing one outcome by correcting for the number of times each button was pressed. Hence, reward and punishment errors on neutral trials were divided by the total number of times each prospective button was pressed before the difference was calculated. A larger value for this measure may suggest higher reward sensitivity. We correlated these two behavioral measures with P2 and N2/EAP, which showed significant contrasts between reward and punishment conditions.

Interestingly, a stronger reward learning bias was correlated with higher P2 difference (*r_s_* = .39, *p* = .018; see Fig. 3), but not N2/EAP (*r_s_* = .18, *p* = .29). In contrast, a larger reward sensitivity was correlated with both higher N2/EAP effect (*r_s_* = .38, *p* = .023), and P2 effect (*r_s_* = .38, *p* = .026).

**Fig. 3.**
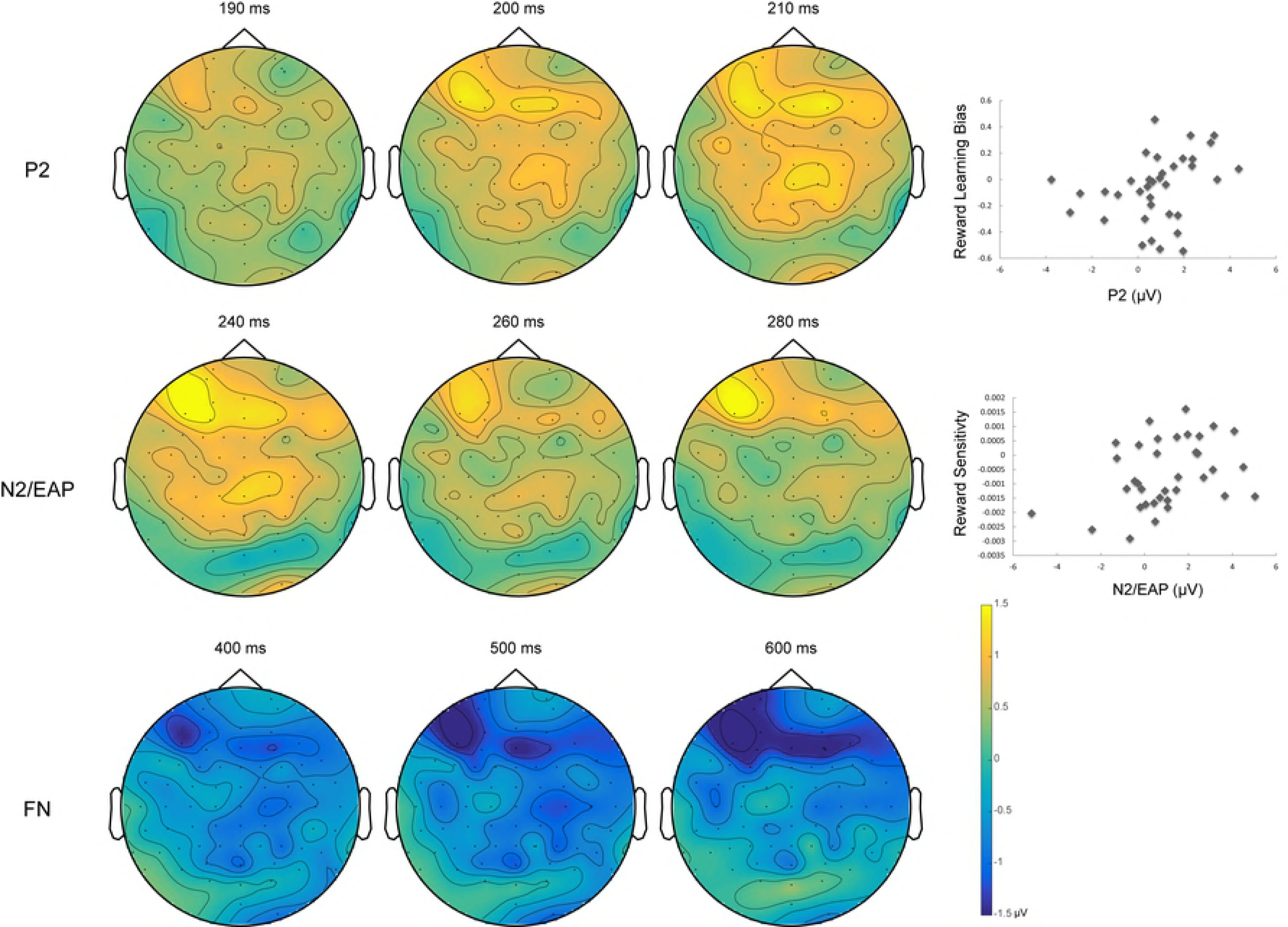
Topographical maps for the three components observed for the current study. All topographical maps are based on difference waves: P2, EAP (reward - punishment), FN (punishment - reward). Note how the N2/EAP and FN do not invert polarity below or between left and right ocular channels, suggesting the source of these activities is neural rather than ocular artifacts. Scatter plots show correlations between P2 contrast and reward bias (punishment errors on reward trials minus reward errors on punishment trials), and between N2/EAP contrast and reward sensitivity (reward minus punishment errors at neutral trials).

## 4. Discussion

In the current study, participants learned to associate uninteresting images with reward, punishment or neutral monetary outcomes. We observed an early bias towards learning reward associations, although all associations were eventually learned. Brain electrophysiological responses to visual presentations of the images during a subsequent implicit task showed differential spatiotemporal effects. Specifically, when images were associated with monetary reward, earlier stages of stimulus detection and evaluation were affected (P2 and N2/EAP; 170 – 310 ms). In contrast, we observed a later post-evaluative effect for images associated with monetary punishment (FN; 350 – 650 ms).

### 4.1. Behavioral effects of associative learning

Even though participants made more attempts towards receiving rewards and avoiding punishments, as indicated by the larger number of reward and punishment button-press than neutral, accuracy and learning trials data suggested that reward associations, but not punishment associations, were preferably learned. One explanation for the different learning profiles is that reward and punishment may act in opposite manner when interacting with behavioral control systems, with reward encourages active approach and punishment facilitates behavioral inhibition [43, 44]. For example, when reward and punishment were orthogonally manipulated with action (button-press versus button-release) in a go/no-go task, Guitart-Masip and colleagues [45] found that reward facilitated learning go responses whereas punishment motivated more no-go responses. Given that our learning task is a go task, our results were consistent with this framework of valence and action interaction. In contrast, when comparing learning and response biases for punishment and neutral conditions, we observed a different profile: punishment cues were learned faster than neutral cues (*p* = .026, uncorrected) and punishment buttons received more preference over neutral buttons. Therefore, we further suggest that the interaction between valence and action seems to influence specific strategies during learning which prompts biases towards reward learning. However, valence (and salience) by itself, positive or negative, also seems to promote learning.

Our accuracy results are in line with previous studies using reward cues to motivate performance, showing that participants are more accurate for high reward trials than low reward or neutral trials [46-48]. We demonstrate here that such reward motivation can be present even when participants simultaneously learned to avoid monetary punishment. However, we did not observe any RT difference reported previously. This could be a result of task instruction: our learning task was self-paced rather than timed, which emphasized accuracy over speed. Therefore, participants may have been motivated to maintain high accuracy rather than faster RT.

### 4.2. Reward associative learning

#### 4.2.1. P2 wave

Our first hypothesis, that the frontocentral P2 elicited by rewarded stimuli would be enhanced was supported. This result is consistent with paradigms using monetary reward to motivate task performance. In such tasks, enhanced P2 to reward cues was interpreted as an indication of motivated stimulus processing to obtain the impending reward [22]. The current result is also consistent with the interpretation that the P2 effect is larger for passively viewed reward-predictive cues due to their increased salience [20, 26], as well as P2 as an index of automatic attentional capture [28]. However, it has been difficult to discern whether P2 was produced by active reward anticipation, learned incentive salience, or attentional capture when EEG recording takes place during reward learning. In the present study, P2 was elicited *after* reward learning during an implicit task in which no form of reinforcement was provided. Therefore, the current study may suggest the possibility that P2 modulations could take place implicitly. Interestingly, making more punishment errors during reward trials cues (i.e., updating reward associations) and less reward errors for punishment cues were correlated with larger P2 effects. Presumably, such an asymmetry further shapes learning and may contribute to later observed P2 difference. Future research utilizing trial to trial analyses could test this possibility.

Other ERP components share a similar frontocentral scalp distribution to the P2, albeit later in time (200-300 ms). These include the feedback-related negativity (FRN; [49, 50]) and the reward positivity (RP; [51]). Both components are elicited by feedback and feedback-predicting cues, with FRN greater for monetary loss and RP greater for monetary gain [52, 53]. It is an ongoing debate whether FRN/RP modulations during this time interval are driven by negative deflections to punishment or positive changes to reward cues [51, 54, 55]. We believe that the P2 effect found in the current study differs from FRN/RP effects reported in previous studies. First, our P2 emerged and peaked earlier (170-230 ms) than the FRN/RP, indicating a temporal dissociation. Additional support for our claim comes from previous functional interpretations of the P2 as stimuli-based attentional modulation whereas FRN was linked to action monitoring or controlled attention [26, 28]. In this regard, the FRN/RP are better associated with the frontocentral N2. This component is thought to play a role in conflict monitoring [51, 56], and produces the baseline for FRN/RP modulations [55]. The frontocentral N2 also is distinguished from the P2 in more traditional visual and auditory attention tasks [57, 58].

We agree with the functional interpretation of P2 and FRN/RP. Importantly, in our recording task participants simply viewed the previously learned cues without any current reinforcement feedback or overt response. It is not surprising that without error feedback or conflict, little activity from the FRN/RP latency was observed in frontocentral sites [59]. The P2, however, might reflect stimulus-specific detection and could remain as long as participants see stimuli previously associated with reward, punishment or neutral outcomes. Interestingly, we did observe activity from the N2 time window, which is discussed in the following section.

#### 4.2.2. N2/EAP

Our second hypothesis that reward cues would show enhanced N2/EAP modulation (210-310 ms) was also supported. The present finding is consistent with previous studies in our laboratory employing appetitive stimuli with heightened incentive salience among certain groups of individuals – such as chocolate and marijuana cues viewed by chocolate cravers and individuals with marijuana dependence [31, 32]. Because such individuals have extensive experience with their craved substance, associated cues are suggested to be attributed with incentive salience as a result of dopaminergic activity [4, 6, 60]. It is also consistent with past studies employing aversive but salient distractors (e.g., spiders, threat words) to capture attention and influence performance [29, 33]. The N2/EAP effect in previous studies has been discussed as an index of automatic attentional capture based on fast visual sampling of significant features [31, 33, 39]. However, salience of the stimuli employed in previous research was likely developed outside of laboratory, either through real-life (marijuana) or evolution (spiders). In the present experiment, we have shown that elevated salience, indexed by the N2/EAP, can be induced with a learning session as short as thirty minutes. Moreover, the N2/EAP (and P2) effect seems to be related to individual differences in reward sensitivity. Individuals who are more reward seeking (making more reward than punishment errors) during neutral trials tend to show larger N2/EAP modulation. Note that reward or punishment errors in this case only inform learning of neutral cues, which did not factor in the observed N2/EAP difference. Given this observation, future research should test individual difference in reward-seeking and punishment avoidance traits and examine how these traits could modulate feedback-related activity during learning and the P2 and N2/EAP after learning.

Unlike the previous study [37], we did not observe any early difference in the occipital site (called very early effects of emotion; VEEE, see Fig. 2). There are two plausible explanations for such discrepancy. First, ERPs in the present study were referenced to the averaged mastoids, whereas VEEE was reported with the average reference, making direct comparison difficult. A second interpretation for the present result stems from the established anatomy of the ascending dopaminergic projections of the reward system, which target frontal cortex locations far more extensively than occipitoparietal ones [61]. This should logically result in more anterior than posterior scalp distribution of reward-dependent electrophysiological activity. Yet, associating reward with highly visually challenging stimuli (i.e., Chinese characters) may selectively enhance early visual processing, establishing a more posterior effect (VEEE; [37]).

#### 4.3. Punishment avoidance learning and FN

We conducted exploratory analyses on an observed frontal ERP effect. A noticeable negative peak with frontopolar distribution (FN) was evident particularly for the punishment condition. A similar FN effect was previously reported in response to failed versus successful inhibitions in a stop-signal task, and was interpreted as reflecting a post-evaluative error correction mechanism [62]. If such interpretation were to apply to our study, a post-evaluative error mechanism may be implicitly activated possibly in the form of punishment associations for trials not requiring an overt categorization or selection and execution of a motor response. Another study reported a similar, albeit later and more sustained frontal negativity (named frontal slow wave; 500 – 1500 ms) during a blackjack task in response to trials associated to losses compared to wins and ties [63]. A frontal slow wave was also reported during a counting Stroop task. Interestingly, such effects were more pronounced among individuals who experienced the greatest reduction in positive affect ratings [64], perhaps suggesting a similar sensitivity of the frontal negativity to the impact of negative affect on cognitive control operations.

Finally, another frontal component argued to be the index of familiarity-driven recognition (FN400; e.g., [65]) showed somewhat similar temporal and spatial distributions as the current FN. It could be that in our study the FN modulation reflected implicit recognition of the learned images. However, in the ERP memory literature FN400 is more *positive* for remembered (old) compared to new stimuli, and this effect is enhanced for images with negative emotional valence [66]. This is at odds with our findings for the FN in the present study, which was more negative in amplitude for punishment relative to reward images. Nevertheless, future research could draw from this exploratory finding, along with previous results, further demonstrating the role of this frontal negativity in the interplay between affect, error correction and cognitive control during punishment avoidance learning.

#### 4.4. Conclusions and limitations

A number of limitations can be identified in the present study. First, our learning point estimation did not take into account potential response biases during interleaved reward and punishment avoidance learning. For example, according to our estimation procedure, a person pressing reward button across all trials would “learn” reward associations at trial one and never learn others. This would not be an accurate estimation of individual’s learning. Future research employing statistical modeling to analyze learning data is needed to address any response bias during interleaved learning (e.g., [67, 68]). Second, we did not collect EEG data during the associative learning task, which prevented us from further analyzing how learning contributes to later differences in perceptual processing. A future extension of the present study could assess the longevity of reward or punishment associations as indexed by the P2, N2/EAP and FN effects reported here. Additionally, although we interpreted our results in relation to associations of reward, punishment and neutral outcomes, we could not rule out effects of perceptual or semantic priming. Indeed, participants may be primed to show differential electrophysiological responses towards our stimuli. Future studies could test this possibility by using the same pictures during learning, and record electrophysiological responses to semantically related words.

In conclusion, using an associative learning task, we found distinct trajectories of learning marked by early preference towards learning reward associations, when there was a need for competition of attentional resources. Despite an absence of behavioral differences among cues at the end of the learning period, we found larger early ERP components (P2 & N2/EAP) for reward over punishment cues. We interpreted them as indices of heightened incentive salience. Additionally, we reported an exploratory finding of a frontal negativity (FN) that appeared to be more prominent for punishment over reward cues. Future research could test the possibility of this frontal negativity involved in mediating how an aversive state can motivate additional needs for control and learning.

### Authors Notes

We thank Megan Liau and research assistants in our lab for the help with data collection. We are particularly grateful to John McDonald and Anne Smith for valuable advice on the project. This research was supported by grants from Natural Sciences and Engineering Research Council of Canada (NSERC) and Canada Foundation for Innovation (CFI) to ML.

## Footnotes

1 IAPS reference numbers: 5740, 7000, 7003, 7004, 7006, 7010, 7012, 7025, 7026, 7031, 7035, 7040, 7080, 7090, 7100, 7160, 7175 and 7182.

